# Risk factors associated with tick infestation and *Ehrlichia ruminantium* amongst cattle in Guadeloupe, French West Indies

**DOI:** 10.1101/2025.08.20.670949

**Authors:** Dufleit Victor, Guerrini Laure, Cozier Joanna, Samut Thony, Pezeron Manuel, et Etter Eric

## Abstract

*Ehrlichia ruminantium* infection and infestations by two tick species, *Amblyomma variegatum* and *Rhipicephalus microplus*, are a major threat to the livestock sector in Guadeloupe. This study examined specific tick infestations for 261 cattle from 73 farms on the archipelago’s three largest islands, Basse-Terre, Grande-Terre and Marie-Galante. A questionnaire was implemented to gather information on farming practices and livestock characteristics in order to identify potential risk factors associated with the presence of antibodies against *E. ruminantium* and the abundance of ticks in livestock. A total of 81% of the inspected cattle were found to be infested with ticks, with a prevalence of 69% for *A. variegatum* and 47% for *R. microplus*. At farm level, 96% of farms had infested animals, 84% of which were infested with *A. variegatum* and 73% with *R. microplus*. Risk factors were first explored using multiple correspondence analysis (MCA) followed by a generalized linear model at the farm level, and finally a generalized linear mixed model at cattle level. MCA revealed that non-traditional breeding system, with free-grazing animals, were associated with serological evidence of *E. ruminantium* circulation and also higher tick infestations. The utilisation of green fodder had a significant impact on the prevalence of seropositive animals and tick abundance. Differences in infestation were observed between the three largest islands of Guadeloupe and have been discussed in this article. The frequency of acaricide bathing and its interaction with acaricide alternation appeared to be significant factors in the mitigation of tick infestations in cattle. However, this study highlights the potential emergence of acaricide resistance within tick populations in Guadeloupe.

## 2 Introduction

Heartwater is a tick-borne disease of ruminants caused by *Ehrlichia ruminantium* (Allsopp, 2015), an obligatory intracellular Gram-negative bacterium. It is classified by the World Organization for Animal Health (WOAH, 2018) as an acute, fatal, non-contagious infectious disease of ruminants. The disease is associated with high mortality rate, especially among ruminant populations naïve to the disease, where mortality can reach 90% (Asamoah and Fatmawati, 2023). Heartwater also poses a considerable economic burden on farmers (Mukhebi et al., 1999). For example, in Zimbabwe, a loss of 5.6 million US $ nationwide was estimated in 1999, and it is considered as the most important tick-borne disease in South Africa according to Spickett et al., 2011.

In Guadeloupe, the presence of *Ehrlichia ruminantium* was first identified in 1980 by Perreau et al. (1980). The pathogen is believed to have been introduced through the importation of western African cattle and ticks during or following the transatlantic slave trade (Beati et al., 2012). The disease is considered enzootic in the territory (Stachurski et al., 2019), with a small number of cases reported annually to the local CIRAD laboratory, the WOAH reference laboratory for heartwater (Rodrigues, 2025 and personal communication). Ticks of the genus *Amblyomma* are recognised as the primary vectors of the disease especially in Africa (Mapham et al., 2017), and *Amblyomma variegatum* being the main vector in the Caribbean region (Camus and Barre, 1995). Another livestock tick, *Rhipicephalus microplus*, which is also widespread in the Caribbean, has recently been identified as a potential vector of *Ehrlichia ruminantium* in western Africa (Biguezoton et al., 2016; Some et al., 2023).

Livestock production in Guadeloupe remains mainly traditional, with tethered breeding being the predominant method for raising cattle, goats and sheep. The sector is currently in decline (Agreste, 2021; Galan et al., 2008) exacerbated by the threat posed to ruminant population by heartwater and its vectors. Furthermore, the costs associated with tick control remain high and constitute a significant burden for farmers (Bernier et al., 2025; Camus, 1989).

A heartwater vaccine is used in South Africa (Van Den Heever et al., 2022) but vaccination is still limited for heartwater because of important genetic diversity (Vachiéry et al., 2008) and limited cross immunity (Vachiéry et al., 2010). No commercial formulation is currently available in the Caribbean. In the absence of effective vaccination and limited prophylaxis, control of tick infestations remains the primary method of preventing the disease. Traditional Creole breeds have previously been reported to show greater resistance to both heartwater and tick infestation (Camus and Barre, 1987). Studies conducted in South Africa, Kenya and Comoros (Mdladla et al., 2016; Peter et al., 2019; Boucher et al., 2020), have investigated potential risk factors associated with *E. ruminantium* seroprevalence. However, no analysis of *E. ruminantium* risk factors have been conducted in the Caribbean. This study presents the results of a survey on tick infestations and associated diseases, as well as risk factors linked with livestock management practices on cattle farms throughout Guadeloupe. Tick infestations in cattle were assessed for the different islands of the Guadeloupe archipelago. A risk factor analysis was performed using data collected from farmers, including management practices and, epidemiological information, to identify key factors associated with the presence and abundance of ticks and *E. ruminantium* seroprevalence (Dufleit et al.2025). This study aims to inform more effective strategies for disease surveillance and control in the region.

## 3 Materials and methods

A cross-sectional survey was conducted to collect cattle serum, ticks for identification and counting as well as data on breeding practices (at farm and cattle levels). The sampling design was previously described in Dufleit et al., 2025.

### 3.1 Farm and cattle Survey

A questionnaire was administered among all cattle farmers, from whom blood and tick samples had been taken, in order to gather information on farming practices and other potential risk factors associated with *E. ruminantium* seroprevalence and tick infestations. A summary of the collected variables is presented in table 1. The questionnaire included questions on whether farmers had received formal agricultural education and whether livestock farming was their primary occupation. Additional data included the number of cattle owned, the daily time spent on breeding activities, and the predominant rearing system (i.e., tethered vs. free grazing). For farmers reporting tethered management, further details were recorded regarding chain length and frequency of stake relocation. The use of green fodder was also documented. Information on tick control practices was gathered, including acaricide use, treatment frequency, date of the last treatment and whether rotation of active substances was implemented. In additions, farmers were asked whether cattle egrets, known to be potential tick disseminators in the region (Corn et al., 1993), were frequently observed in proximity to their livestock.

**Table 1.** Summary of collected potential risk factors at the farm and cattle levels (^*^ indicates risk factor collected at the farm level but also included in analysis at the cattle level)

| Question | Variable Name |
| --- | --- |
| Factor collected at farm level |  |
| Island where breeding is located | island |
| Agricultural education | agriFormation |
| Number of cattle owned | EfTot |
| Breeding is main activity | pActivity |
| Average daily time spent on breeding | workTime |
| Are your animals tethered | stakeBreeding* |
| Length of animal's tether chain | chainLength* |
| Frequency of movement of cattle stakes | stakeMvmtFreq* |
| Use of green forage | greenForage* |
| Frequency of acaricide use | acarFreq* |
| Alternance of active substance | acarAlternance* |
| Observation of cattle egret around animal | cattleEgret |
| Factor collected at cattle level |  |
| Age of the animal | age |
| Sex of the animal | sex |
| Is the animal gestating | gesta |
| Phenotype of the animal | phenotype |
| Delay in days since last acaricide treatment | lastAcar |

Cattle breed was estimated based on farmer’s declarations, confirmed by phenotype observations and classified as either “creole” (the traditional local breed in Guadeloupe) or “crossbred”. Age (in months) was determined from official identification cards, or alternatively through farmer’s declarations. Additional animal level data included sex and, for female, gestation status.

### 3.2 Hazards identification

#### 3.2.1 Tick infestations

Tick infestation assessments were conducted for the two primary species: *Amblyomma variegatum* and *Rhipicephalus microplus*. Tick species, sex and life stage were identified through palpation and visual examination of cattle body, following the methodology described by Stachurski (2000).

Tick counts at the individual level were transformed into categorical variables using rules inspired by (Bianchi et al. 2003):

– Tick count = 0 → categorical class: 0
– Tick count ∈ [1; 10] → categorical class: 1
– Tick count ∈ [11; 50] → categorical class: 2
– Tick count > 50 → categorical class: 3

Three indicators of infestation per animal were then constructed:

1. *A. variegatum* infestation indicator based on the total count of *A. variegatum* (males, females, nymphs, and larvae).
2. *R. microplus* infestation indicator based on the total count of *R. microplus* (adults and nymphs; no larvae were observed).
3. Overall infestation indicator derived from the summation of the number of ticks of each of the two species.

Mean tick infestation rates (MTIR) were then computed at the farm level, and farms were categorized into three infestation classes, for each of the two tick species and for the overall infestations:

– Low infestation rate: MTIR ∈ [0; 10]
– Moderate infestation rate: MTIR ∈ [11; 50]
– High infestation rate: MTIR >50

Finally, two binary variables were created at farm level to indicate the presence or absence of *A. variegatum* and *R. microplus*, respectively. A final categorical variable named “infestations type” was generated with four classes: “No infestations”, “only *A. variegatum*”, “only *R. microplus*” and “both species”.

#### 3.2.2 Ehrlichia ruminantium serology

ELISA Map1B (Mondry et al., 1998; Semu et al., 2001) were used to assess the serological status of sampled cattle for *Ehrlichia ruminantium*. Detailed laboratory procedures were provided by Dufleit et al., 2025. Tests results were aggregated at the farm level: if at least one animal from a farm tested positive with the ELISA Map1B, the farm was classified as positive.

### 3.3 Statistical analysis

#### 3.3.1 Bivariate analysis

As a preliminary step, a chi-squared test was performed at the farm level to detect significant differences between *E. ruminantium* serological status and tick infestation classes. *E. ruminantium* serological status and tick infestation classes were then analysed separately.

#### 3.3.2 Multiple Correspondence analysis (MCA)

Questionnaire data were cleaned and compiled into a csv file for analysis using the R programming language (R Core Team, 2023). Numerical variables were discretized into two categories using the median. For the variable “workTime”, the third quartile was preferred to differentiate farmers that spend most of their time on their farm from others. The relationships between potential risk factors and both, serological status to *E. ruminantium*, and tick infestation levels were analysed using Multiple Correspondence Analysis (MCA; Mori et al., 2016) implemented via the FactoMineR R package (Lê et al., 2008). Risk factors assessment for *E. ruminantium* seropositivity was only done at farm level as the WOAH country manual statement (WOAH, 2018) stated that the Map1B ELISA should be considered at the herd level only, not at the individual level. In order to assess the risk factors for tick infestations, MCA were carried out for the global tick infestations, for *A. variegatum* and for *R. microplus* infestations at both farm and animal level. A maximum of ten dimensions were extruded for each analysis. Dimensions were selected for graphical representation with the selection being made on the basis of the highest correlation ratio (eta-squared) found for the variables of interest (*E. ruminantium* status or tick infestations). Missing values (NAs) were managed using the ‘completeMCA’ function from the *missMDA* R package (Josse and Husson, 2016).

#### 3.3.3 Generalized Linear Models (GLM) and Generalized Linear Mixed Models (GLMM)

Variable selection for modelling was primarily guided by the factor plots from the preceding MCA. Variables positioned near, opposing, or forming a structure with the variable of interest in the MCA space were considered relevant. A Generalized Linear Model (GLM) (Dobson, 2002) was employed to study heartwater serology and mean tick infestations (*A. variegatum* and *R. microplus*) at farm level. To analyse tick infestation at the cattle level, a Generalized Linear Mixed Model (GLMM) (Bolker et al., 2009) was preferred due to its ability to account for clustered data, considering the sampling of several animals per farm. In this model, the farm identifier (EDE) was included as a random effect to account for the grouping structure. A binomial link function was used for heartwater serology analysis, while a Poisson link function was applied for the tick infestation models. The lme4 R package (Bates et al., 2015) was used to implement the GLMM.

To further investigate species-specific associations, the “infestations type” variables (with four classes) were included in the serological model. This allowed to assess whether farms with seropositive animals were more likely to be associated with one tick species rather than the other.

At the farm level, infestation indicators were calculated as the average tick counts per farm; these averages were rounded to the nearest integer for compatibility with the Poisson distribution assumed by the GLM. For all analysis, continuous variables (e.g., chain Length, stake change time, acaricide treatment frequency) were included as numerical predictors and were not discretized, unlike in the preceding MCA.

In cases where predictors were interdependent (e.g., chain length and stake movement frequency being conditional on the animals being tethered), the tethered vs. free grazing variable was preferred during model construction.

Potential interactions between predictors were also considered according to previous study (Salas et al., 1986):

– acaricide treatment frequency, alternance of active substances and herd size interactions;
– herd size and Island interactions.

To detect multicollinearity, Variance Inflation Factors (VIF) were calculated using the *car* R package (Fox and Weisberg, 2019): when VIF exceeded a score of 5, collinearity between variables were compared to select the most informative predictor (Zuur et al., 2010).

Model optimizations were performed using backward stepwise methodology using the ‘step’ function from the R stats package (R Core Team, 2023) for GLM and the ‘GLMERselect’ function from the *StatisticalModels* (Newbold, 2023) package for GLMM. Odds ratio (OR) (Kerr et al., 2023) for binomial GLM and incidence rate ratios (IRR) for Poisson GLM and GLMM with 95% confidence intervals were computed. At the cattle level, two models were constructed, one including the delay since last acaricide treatment as predictor and one excluding this variable to better overview the effects of other risk factors on tick infestations (Baguley, 2018). Finally, a goodness of fit procedure was conducted by comparing selected models with their null counterparts using one-way analysis of variance based on the chi-squared test (Coxe et al., 2009).

## 4 Results

### 4.1 Descriptive analysis

#### 4.1.1 *Ehrlichia ruminantium* serology and tick infestations

A total of 73 farmers responded to the questionnaire. Blood samples and tick counts were collected from one to six cattle per farm, resulting in a sample of 261 cattle. The 76 animals tested positive for *E. ruminantium* (28% [22% - 34%]_95%_) identified 47 farms (63%) with at least one MAP1B ELISA positive animal (Dufleit et al., 2025). Cattle tick infestation counts ranged from 0 to 241 ticks per animal (0 to 150 for *A. variegatum* and 0 to 240 for *R. microplus*) with a median/mean infestation of 10/32.15 ticks (3/12.61 for *A. variegatum* and 0/19.54 for *R. microplus*). Regarding the classification of tick infestation, 49 cattle (81 for Av, 138 for Rm) were found to be entirely free of ticks, 84 (103 for Av, 47 for Rm) exhibited 1 to 10 ticks, 74 (63 for Av, 42 for Rm) exhibited 11 to 50 ticks and 54 cattle (14 for Av, 34 for Rm) exhibited more than 50 ticks. According to these results, the prevalence rates of infestation were 81%, 69% and 47% for all ticks, *A. variegatum* and *R. microplus* respectively.

At farm level, MTIR results showed a low infestation within 31 farms (46 farms for *A. variegatum* and 45 farms for *R. microplus*), a moderate infestation within 27 farms (22 farms for *A. variegatum*, 21 farms for *R. microplus*) and a high infestation within 15 farms (5 farms for *A. variegatum*, 7 farms for *R. microplus*). Only 5 farms were found to have non-infested animals, 12 were free of *A. variegatum* and 19 farms had no observation of *R. microplus*. Thus, the prevalence rate of tick infestation at the farm level was 0.96, 0.84 and 0.73 for all ticks, *A. variegatum* and *R. microplus* respectively.

Tick distribution on the Guadeloupean territory at communal scale showed the presence of both tick’s species in all the investigated municipalities except for Saint-Claude and Les Abymes where only *R. microplus* were found (Fig.1).

**Figure 1.**
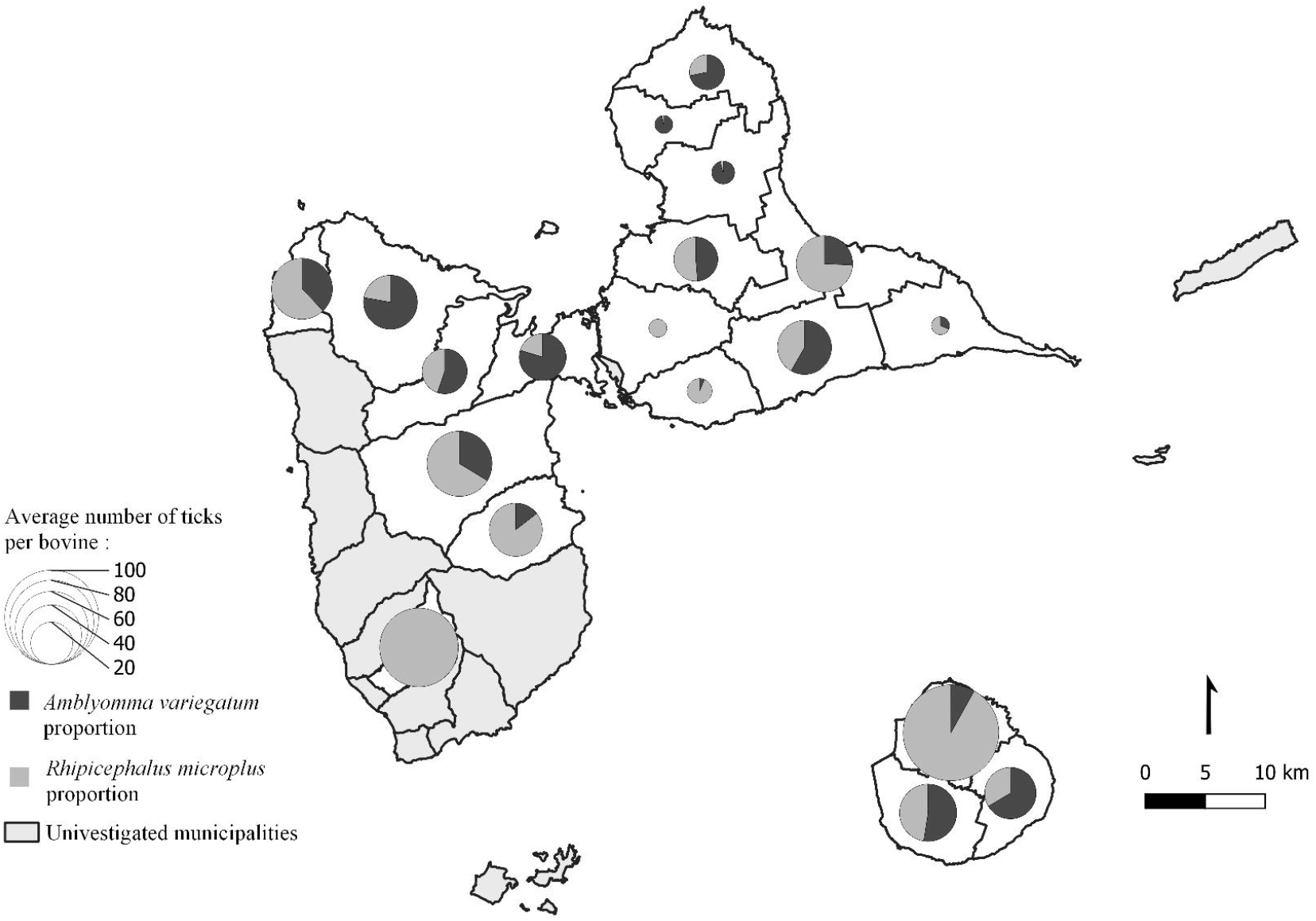
spatial distribution of mean cattle tick infestations over municipalities of Guadeloupe.

No significant statistical association between *E. ruminantium* serological status and overall tick infestation class were found at the farm level (Table 2; p-value=0.056). Serological status and tick infestations were analysed separately in the multivariate analyses.

**Table 2.** *E. ruminantium* serological results at farm level according to tick infestation classes.

| Tick Infestation Class | Number of farms with *E. ruminantium* seropositive cattle | Number of farms with no *E. ruminantium* seropositive cattle |
| --- | --- | --- |
| +50Tick | 9 | 6 |
| 11-50Tick | 22 | 15 |
| 0-10Tick | 16 | 15 |

#### 4.1.2 Questionnaire responses and cattle’s descriptions

Among the 73 farmers who completed the questionnaire, 15 (20%) were from Basse-Terre, 39 (54%) from Grande-Terre and 19 (26%) from Marie-Galante. The predominant activity among 33 (45%) respondents was cattle breeding, and 23 (32%) reported having received an agricultural education during their schooling. The median time spent on animal husbandry was 3.5 hours per day, with the third quartile (5 hours) demarcating the upper limit for the creation of two categories: 16 breeders (22%) spending more than 5 hours daily to breeding activities. Regarding management practices, 66 breeders (90%) tethered their animal, with a median chain length of 8 metres, and a median frequency of stake movement of 1.5 days. Furthermore, 16 breeders (22%) provided green fodder to their cattle, while 62 (85%) observed the presence of cattle egret around their herds. The median acaricide application frequency was twice per month, and 20 breeders (28%) employed a rotation of active substances when treating their animals. The median delay since the last acaricide treatment was 15 days.

From the 261 cattle sampled, 56 were selected from Basse-Terre, 137 from Grande-Terre, and 68 from Marie-Galante. Among these animals, 209 (80%) were females, and 52 (20%) males. At the time of sampling, 83 females (40%) were pregnant. The median age of the animals was 48 months, with 133 cattle under 4 years old, and 128 over 4 years old. The breeds included 150 Creole animals and 111 crossbred animals.

### 4.2 Risk factor analysis of *Ehrlichia ruminantium* seroprevalence

In the MCA assessing risk factors associated with *E. ruminantium* seroprevalence, the first two dimensions, explaining 18% and 15% of the total dataset variance respectively, were selected as they provided also the most informative representation of *E. ruminantium* serology variables. These dimensions had eta2 values of 0.18 and 0.37 respectively for the *E. ruminantium* variable (fig.2).

Observation of variables graph obtained using the MCA revealed several factor associations. The *E. ruminantium* serological results variable showed similar categorical structure with the tick presence variables for both tick species. Absence of serological evidence of *E. ruminantium* showed proximity with both categories accounting for absence of *A. variegatum* and *R. microplus* respectively. The two tick presence categories showed also proximity with the *E. ruminantium* serological evidence category. The “free grazing” variable was also closely associated with the *E. ruminantium* serological evidence category. Farms with more than seven animals appeared to be associated with positive serological results on the farm. High-frequency treatment was strongly related to the presence of serological evidence, while the opposite was observed for low-frequency treatment. The three variables “workTime”, ‘pActivity’ and “agriFormation” exhibited spatial proximity with the acaricide frequency variable. Moreover, low treatment frequency was associated with farmers who worked more than 5 hours a day on their livestock, for whom livestock farming was the main activity, and who had received agricultural training at school. The use of green fodder and the alternation of acaricides and active substances were significantly represented on the second dimension and appeared to be associated with the presence of serological evidence of *E. ruminantium*. The absence of observations of bovine egret showed proximity to the absence of serological evidence of *E. ruminantium*.

Variable selection for the GLM was guided by the descriptive analysis of the MCA variable category plot (fig. 2). The selected variables included the presence of *A. variegatum* and *R. microplus* ticks. Other retained factors were farming system (livestock on stakes vs. free grazing), acaricide frequency and alternance, total effective of cattle in the farm, observations of cattle egrets and the use of green fodder.

**Figure 2.**
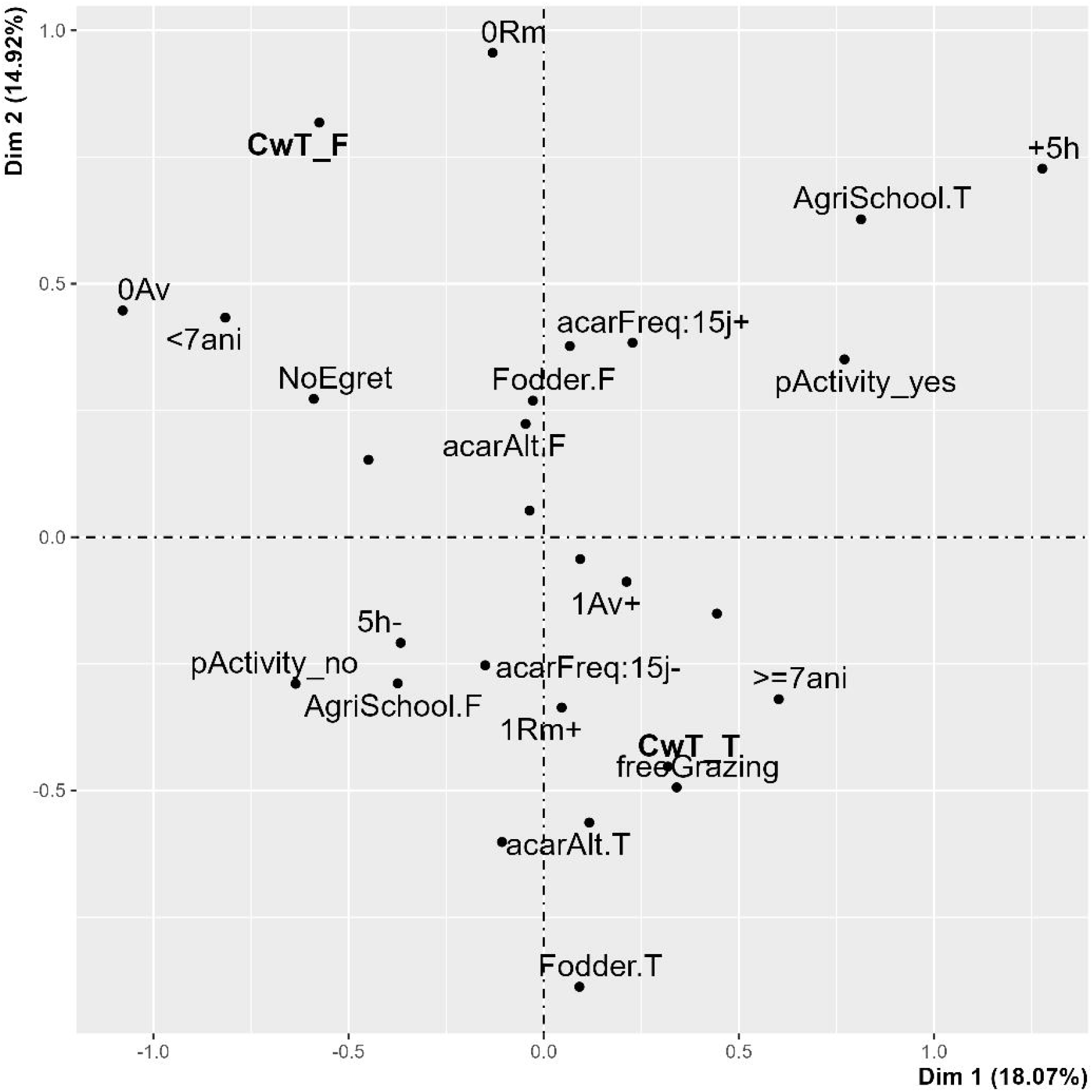
Multiple correspondence analysis map of potential risk factors associated with *E. ruminantium* seroprevalence (graph of variables categories). Only factors showing the strongest interactions with *E. ruminantium* serological results are labelled.

The variables selected as potential risk factors through the MCA were included in the binomial GLM as explanatory variables for heartwater positivity at the farm level. Finally, the “green fodder” was the only variable showing a statistically significant association with *E. ruminantium* status of farms (Table 3). Specifically, the use of green fodder increased the odds of *E. ruminantium* status by a factor of 5.75.

**Table 3.** Summary of GLM models obtained for Ehrlichia ruminantium seropositivity at the farm level.

|  |  |  |  |
| --- | --- | --- | --- |
| Ehrlichia ruminantium farm status: |  |  |  |
| Family = Binomial |  |  |  |
| Variable name |  | OR ; [Confidence Interval]<sub>95%</sub> | p-value |
| Use of green fodder | No | ref |  |
|  | Yes | 5.75 ; [1.43-38.84]<sub>95%</sub> | 0.03* |

### 4.3 Risk factor analysis of tick infestation

In the MCA exploring risk factors associated with tick infestation at the farm level, the second and fifth dimensions were selected for graphical interpretation. They accounted for 13.69% and 7.8% of the total dataset variance, respectively (fig. 3). The MTIR variable had eta2 values of 0.29 for the second dimension (x-axis) and 0.32 for the fifth dimension (y-axis), representing the maximum values found across the ten extruded dimensions. The MTIR increased along the second principal dimension (x-axis) with a negative value for the low infestations category and increasing positive values for the medium and high infestations categories. Several variables were spatially associated with the low infestation category, including: the use of green fodder, farmers spending more than 5h hour per day on their exploitation and the absence of acaricides alternation variables. Conversely, alternation of acaricides showed proximity with the two highest infestation categories. The island factor exhibited specific proximities with Grande-Terre showing proximity with the low infestations category and Basse-Terre showing proximity with the highest MTIR category. Looking specifically at the second dimension (projection on the x-axis) Marie-Galante is also on the same side as the highest MTIR category. Similar results were observed with the observations of cattle egret and the “livestock on stakes / free grazing” variables: “no egret” and “free grazing” categories were associated with highest MTIR category while “stakesBreeding” category was associated with low tick infestation rates.

**Figure 3.**
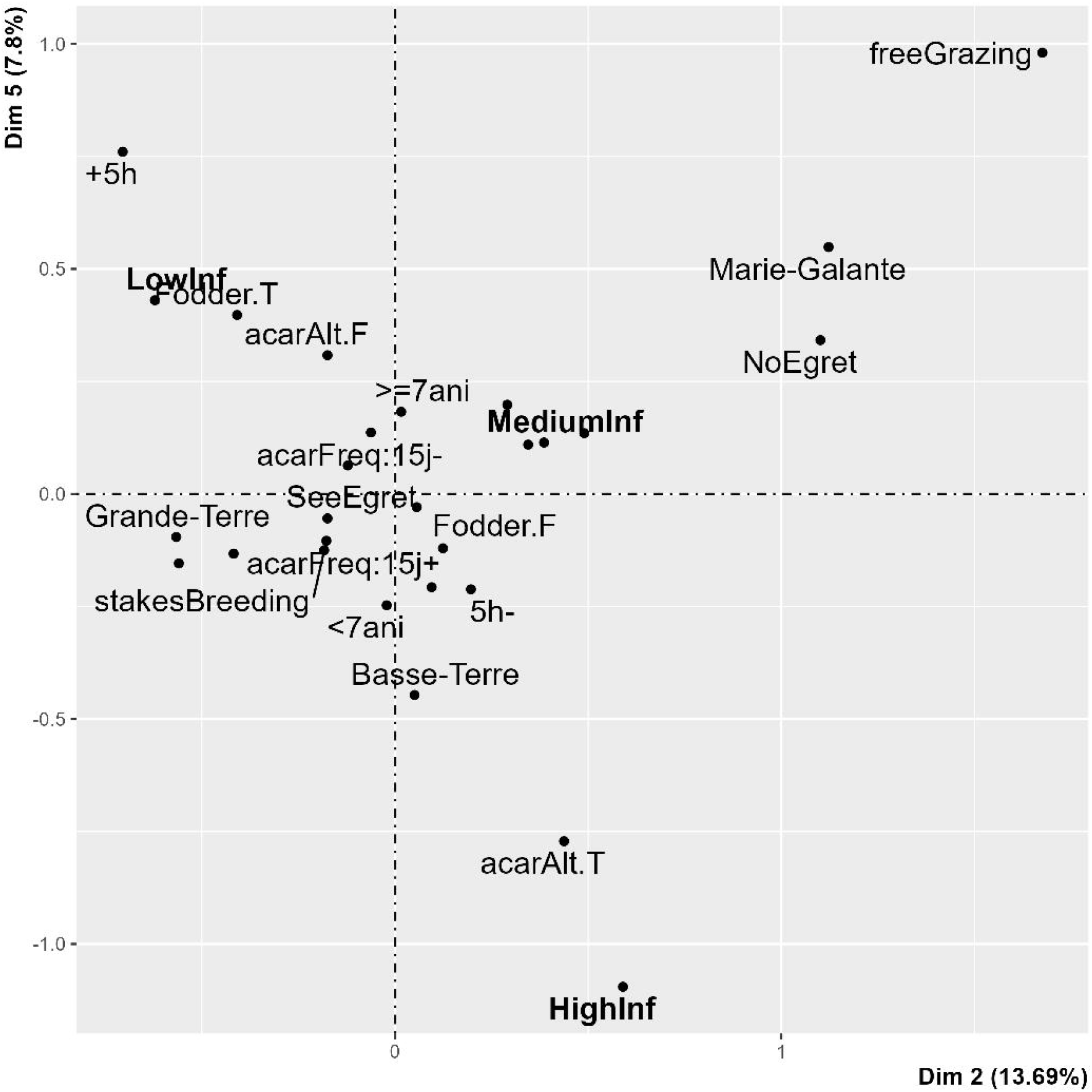
Multiple correspondence analysis map of potential risk factors associated with MTIR at farm level (graph of variables categories). Factors with the highest interactions with MTIR are labelled.

In the MCA of tick infestation at cattle level, the first and fourth dimensions were retained, with eta2 values for tick infestation indicator of 0.27 and 0.42, respectively. These dimensions explained 13.26% (dimension 1) and 8.58% (dimension 4) of the total variance of the dataset (fig.4). As for the analysis at farm level, the tick infestation indicator is increasing along the x-axis (first dimension). Grande-Terre was associated with the lowest infestations categories, and Basse-Terre and Marie-Galante showed proximity with the highest infestation categories. The three variables accounting for use of acaricides showed also clear structure regarding tick infestation. Acaricides alternation, the categories “acaricides frequency > 15 days” and “last acaricide treatment > 15days” were associated with the highest infestations categories. Their alternate categories were associated with low tick infestations categories. Creole cattle also appear to be more infested than crossbred one, with high proximity of the crossbred variables with the no tick infestations class. The age of animals also showed interactions with infestations categories, “older animals” category appeared closer to high infestation categories while “younger animals” category was closer to low infestation categories.

**Figure 4.**
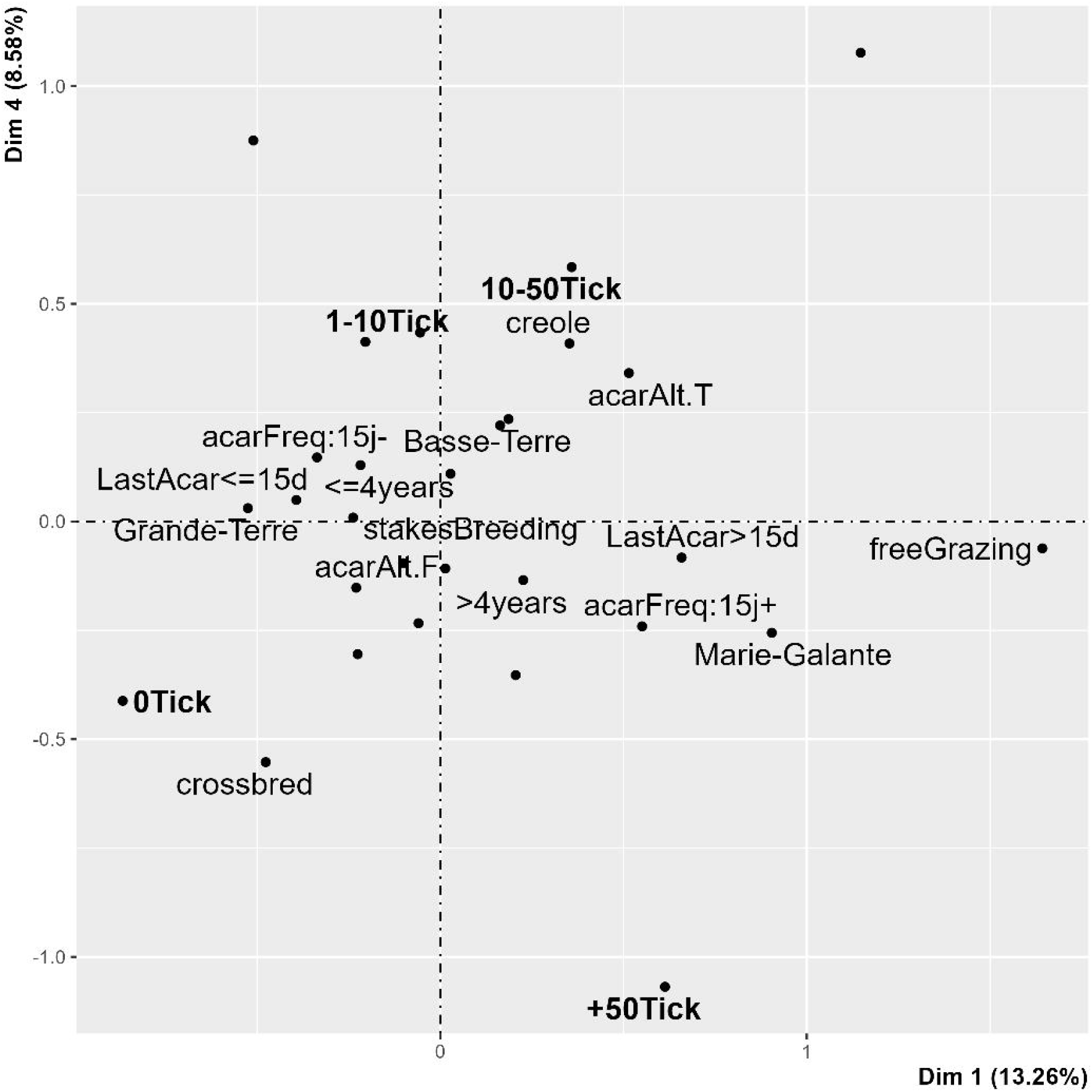
Multiple correspondence analysis map of potential risk factors associated with tick infestation indicator at cattle level (graph of variables categories).

Based on the descriptive analysis of the factors graph, the following variables were selected for the GLM at the farm level: island, acaricide frequency and alternation, number of animals, farming system (livestock on stakes vs. free grazing), cattle egret observations, use of green fodder, and time spent in the farm.

At the cattle level, the GLMM included the following predictors: time since the last acaricide treatment, island, acaricide frequency and alternation of acaricide use, breeding system (livestock on stakes vs. free grazing) and cattle phenotype.

At the farm level, GLM results (table 4) indicated a significant negative association between the variable “acaricide frequency” and the mean tick infestation rate, with an IRR of 0.53 [0.47 - 0.6]_95%_. Alternance of active substances of acaricide presented a significant positive effect on MTIR with an IRR of 2.96 [2.43 – 3.61]_95%_. In addition, the two variables interacted significantly. Daily time spent on the farm was significantly negatively associated with MTIR, with an IRR of 0.98 [0.96 - 1.00]_95%_. The use of green fodder also showed a statistically significant protective effect, with an IRR of 0.66 [0.57 - 0.75]_95%_. The number of cattle owned did not have significant effect on MTIR (p-value > 0.05). Concerning the island variable, Marie-Galante was found to have the highest MTIR with significant IRR of 0.14 [0.1; 0.19]_95%_ and 0.16 [0.11;0.22]_95%_ for Grande-Terre and Basse-Terre respectively (p-value >0.05). Additionally, the island variable had significant interactions with acaricide frequency and cattle herd size variables.

**Table 4.**
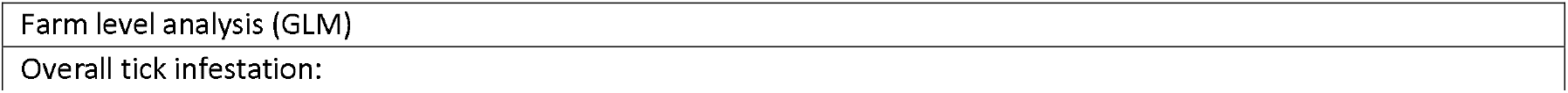
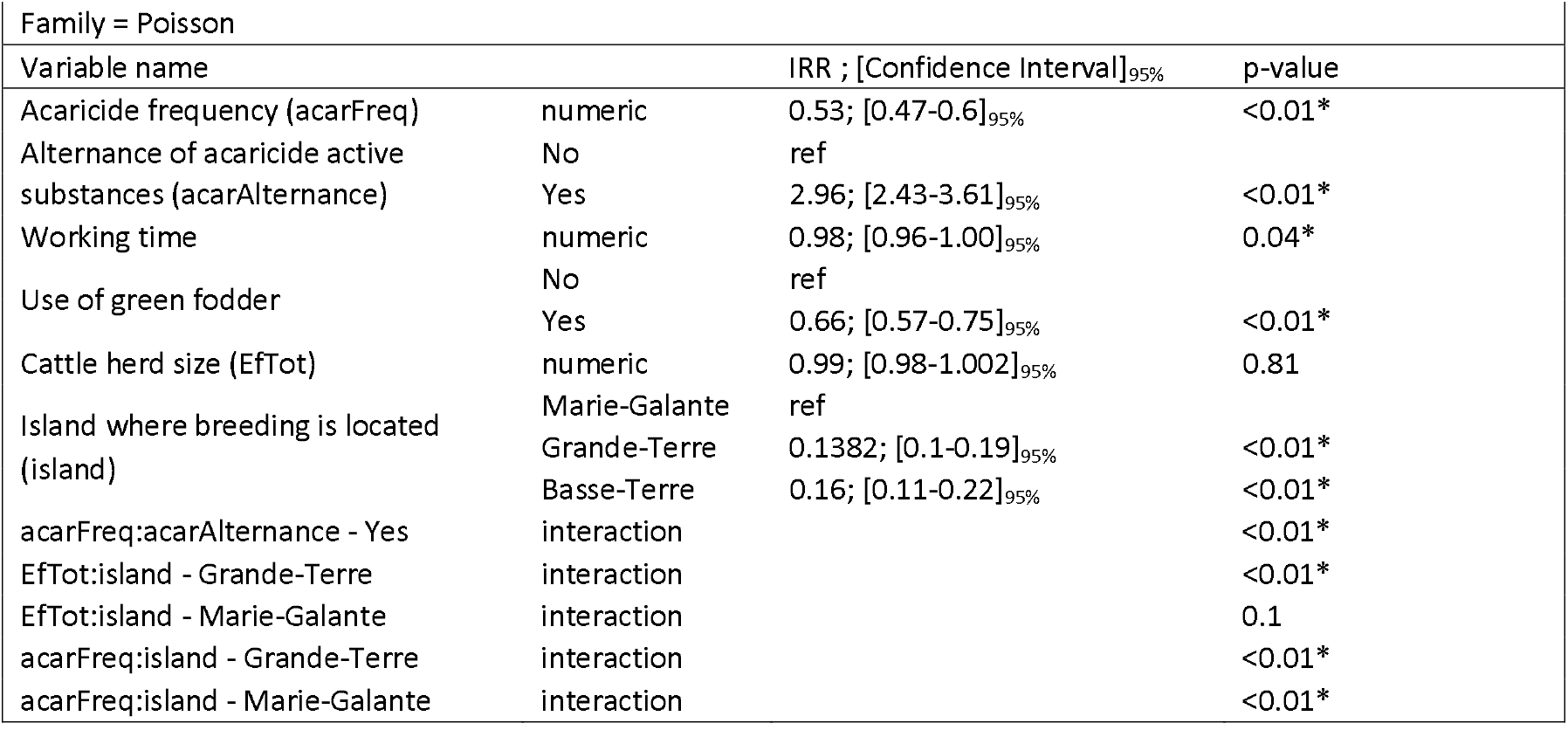
Summary of GLM results for overall tick infestations at farm level.

**Table 5.** Summary of GLMM results for overall tick infestations at the cattle level.

| Cattle level analysis (GLMM) |  |  |  |
| --- | --- | --- | --- |
| Overall tick infestation: |  |  |  |
| Family = Poisson |  |  |  |
| Variable name |  | IRR ; [Confidence Interval]<sub>95%</sub> | p-value |
| Delay since last acaricide treatment (DelayTrt) | numeric | 1.03; [1.02 - 1.05]<sub>95%</sub> | <0.01* |
| Alternance of acaricide active substances (acarAlternance) | No | ref |  |
|  | Yes |  | 0.09 |
| DelayTrt:acarAlternance | interaction |  | <0.01* |
| Overall tick infestation: |  |  |  |
| Family = Poisson |  |  |  |
| Variable name |  | IRR ; [Confidence Interval]<sub>95%</sub> | p-value |
| Island where breeding is located (island) | Marie-Galante | ref |  |
|  | Grande-Terre | 0.27; [0.12 - 0.68]<sub>95%</sub> | 0.01* |
|  | Basse-Terre |  | 0.64 |
| Cattle phenotype | Creole | ref |  |
|  | Crossbred | 0.81; [0.76 - 0.86]<sub>95%</sub> | <0.01* |

At the cattle level, a first model was fitted including the variable delay since last acaricide treatment (in days) (table 4). This variable was the only significant predictor, positively associated with tick infestation (IRR =1.03 [1.02 - 1.05]_95%_). The interaction between “delay since last acaricide” and the acaricides alternation was also significant while “acaricides alternation” itself was not significant. In the second model, excluding the variable “delay since last acaricide treatment”, cattle from Grande-Terre were found to be significantly less infested than those from Marie-Galante (IRR = 0.327 [0.12 - 0.68]_95%_. Cattle from Marie Galante did not show a significant difference in infestation compared to Marie-Galante. Crossbred animals appeared to be significantly less infested by ticks than creole cattle with an IRR of 0.81 [0.76 – 0.86]_95%_.

All final models presented above showed significant differences when compared to their null counterparts. All risk factors showed low collinearity with VIF index below 3 for all variables of all final models. Detailed information regarding specific infestations by *Amblyomma variegatum* and *Rhipicephalus microplus* is provided in the supplementary materials (appendix 1-5).

## 5 Discussion

This study presented results of tick infestations on cattle farms in the Guadeloupe archipelago and investigated the risk factors associated with *Ehrlichia ruminantium* seroprevalence in cattle and tick infestations. Overall, 93% of farms and 81% of cattle surveyed were found to be infested with ticks. These infestation rates are notably higher than those reported in earlier studies: in 2003 and 2005, tick infestation prevalence at the farm level was 35.6% in Guadeloupe and 73.1% in Marie-Galante, respectively (Molia et al., 2008; Vachiéry et al., 2008). Interestingly, the farm typology described by Salas et al. (1986) continues to reflect the current situation. Most respondents to our questionnaire reported engaging in several economic activities, with livestock rearing being only a complementary/leisure activity. Stake rearing remains the dominant livestock management system, while free-grazing systems are still limited to a minority of farms. These farms were mainly located in the south of Basse-Terre in this study. At the individual animal level, historical infestation rates were 14% in Guadeloupe in 2003 and 42.3% in Marie-Galante in 2005. In contrast, our study revealed significantly higher prevalence rates: 90% of farms and 76% of cattle were infested in mainland Guadeloupe (Grande-Terre and Basse-Terre), while in Marie-Galante, 94% of cattle were infested, and ticks were found on all farms visited (100%). These results suggest a marked increase in tick infestation over the past two decades. This discrepancy may be attributed to the timing of the 2003 and 2005 surveys, which were conducted during the implementation of the Caribbean Amblyomma eradication Program (CAP; Pegram et al., 1998; Pegram and Eddy, 2003), with the participation of the French Caribbean region. In Guadeloupe, a limited number of sanitary defence groups called “*Groupements de Défense Sanitaire*” (GDS) (Barré et al., 1996) were created and involved in tick control, through the regular application of acaricide. Since the end of the CAP (Pegram et al., 2004, 2007), all the GDSs in Guadeloupe have ceased operations, which may account for the observed resurgence of tick infestations. The establishment of a new GDS, SANIGWA (Insee, 2020), in 2020, represents a first milestone for a better transfer of knowledge from research to farmers on the territory, in the fight against parasitism.

### 5.1 Risk factors of Ehrlichia ruminantium

Our study identified the use of green fodder as a potential risk factor for *E. ruminantium* seropositivity, as supported by both MCA and GLM analysis. A plausible explanation is that fodder-fed are generally less mobile, potentially increasing their cumulative exposure time to ticks. Additionally, the transport of green fodder may facilitate ticks introduction into farms, as already suggested by Mureithi and Mukiria (2015). However, further research is required to clarify the relationship between fodder-based husbandry practices and *E. ruminantium* transmission. Interestingly, no significant association was observed between the presence or abundance of ticks and *Ehrlichia ruminantium* seropositivity. The observed serology is not the result of the observed infected tick infestation, as MAP1B antibodies can persist for up to 6 months in cattle (Boucher et al., 2020; Semu et al., 2001). While Mdladla et al., 2016 reported an increased risk of *E. ruminantium* seropositivity in the presence of tick vectors, the high overall prevalence of tick infestation in our dataset resulted in a low number of “tick-free” farms, possibly limiting the detection of such associations. The MCA (fig. 2) conducted on *E. ruminantium* serology revealed a potential association with the presence of the two tick species *Amblyomma variegatum* and *Rhipicephalus microplus*. This finding could reflect a common typology of suitable habitat for these two tick species. However, the contribution of *Rhipicephalus microplus* to the epidemiology of *E. ruminantium* needs to be further evaluated (Some et al., 2023). The MCA graph also revealed a strong association between farms with seropositive animals and the free grazing management system. This suggests that free-ranging cattle are exposed to a wider range of habitats, and, consequently, a greater number of potentially infected ticks.

Moreover, we observed associations between the acaricide frequency variable and the three variables, “main activity”, “working time” and “agricultural training”. The existence of these associations could be indicative of a correlation between professional engagement in agriculture and the utilisation of acaricides for cattle. This professional demographic, characterised by agricultural training, full-time dedication, and extended working hours (although this profile is infrequent within the sample population, as previously observed in Guadeloupe (Salas et al., 1986)), exhibits a tendency to employ acaricides on a biweekly basis or less frequently. This low frequency of treatment might be explained by the use of flumethrin, a very effective and persistent acaricide substance, nevertheless this is also the most expensive product in Guadeloupe (Barré et al., 1995)). A better knowledge of management and acaricides, acquired through a specialised scholarship, could also explain lower frequency of treatment among these farmers. However, these potential risk factors showed no explicit relation with the *Ehrlichia ruminantium* serology variable. In addition, the acaricide frequency variable had no significant effect with the GLM models.

The analysis of *E. ruminantium* serology risk factors was conducted exclusively at the farm level, due to the specificity of the test used (WOAH, 2018). Previous studies in Africa and China have investigated the prevalence of *E. ruminantium* in bovine blood samples using molecular detection of the pCS20 gene (Esemu et al., 2018; Guo et al., 2018; Matos et al., 2019). Combining serological data with molecular detection would provide a more comprehensive understanding of the epidemiology and dynamics of *E. ruminantium* infections at the individual animal level. However, due to the short duration of *E. ruminantium* bacteraemia (Camus, 1989), the collection of valuable samples would require a syndromic surveillance system involving farmers and veterinary services for early detection. Such a network previously existed in Guadeloupe through the RESPANG project (Laurent et al., 2012) but is no longer operational. Reactivating expert surveillance systems like RESPANG, could be crucial for improving our understanding of *E. ruminantium* epidemiology and its associated risk factors.

### 5.2 Risk factors of tick infestations

At the farm level, several risk factors were significantly associated with the abundance of ticks on cattle. Increasing the frequency of acaricide applications showed a statistically significant association with reduced tick infestations (table 4), suggesting effective control by acaricide treatments. We obtained similar results when considering *Amblyomma variegatum* infestations separately, but no significant effect of acaricide frequency was detected for *Rhipicephalus microplus* infestations alone (cf. supplementary materials). This last observation could indicate the presence of acaricide resistance among in *R. microplus* populations (Bernier et al., 2025). The greater ability of *R. microplus* to develop resistance (Rodriguez-Vivas et al., 2018), compared to *A. variegatum*, may be explained by its short, monoxenic life cycle. The trophic specialization of *R. microplus* for livestock (Jain et al., 2020), particularly for the immature stage compared to *A. variegatum*, could be linked to a higher rate of exposure to acaricidal substances at population level, potentially increasing rates of resistance development. Interestingly, the alternation of active substances, another important variable was positively associated with high MTIR. Although acaricide alternation is a well-established practice to delay the development of acaricide resistance (Hüe, 2022), this result may reflect reactive rather than proactive management practices. The use of green fodder was associated with a significant lower tick infestation by both MCA and GLM analysis. One hypothesis is that grass cutting, if practised around cattle, would effectively reduce the carrying capacity of the pasture for the tick population and limits the presence of small mammals (e.g. mongooses, rodents), that act as alternate hosts (Barre et al., 1988). By reducing these alternative host populations through vegetation management, the overall tick burden may be mitigated (Fabbro, 2015). The “island” variable strongly influenced tick infestation levels. Farms of Marie-Galante exhibited substantially higher tick infestation levels compared to the other islands. This disparity could be attributed to socio-economic constraints: Marie-Galante has lower average incomes, more limited agricultural infrastructures, and faces higher acaricide prices due to its double insularity, leading to more restrained use of tick control products (Reif et al., 2022). All these risk factors were highlighted within the GLM analysis that accounted for significant interactions between “island” and “acaricide frequency” as well as between “island” and cattle count. Species-specific infestation trends further highlight ecological differences: *A. variegatum* infestations were significantly lower on Grande-Terre compared to Basse-Terre (appendix 5), likely due to Grande-Terre’s drier climate, which is less suitable for *A. variegatum* (Estrada-Peña et al., 2007; Stachurski, 2000). Conversely, *R. microplus* infestations were higher on Grande-Terre (appendix 5), consistent with its known tolerance for drier environments (EstradaLPeña et al., 2005). Additionally, when accounting for the interactions between the island factor and herd size at farm level, herd size showed a positive association with *R. microplus* abundance (appendix 5), possibly reflecting management difficulties in larger herds.

At the cattle level, the most significant risk factor associated with tick infestation was the time since the last acaricide treatment. These results demonstrated the effectiveness of acaricide treatment, as tick counts increased progressively with longer intervals following treatment. Nevertheless, in some specific cases, ticks were still observed on cattle despite recent acaricide application (within 1-3 days), with infestation levels reaching up to 92 ticks per animal. This observation suggests the potential presence of acaricide resistance in tick populations in Guadeloupe (Bernier et al., 2025). It is noteworthy that the time elapsed since last acaricide treatment did not emerge as a significant risk factor when considering *A. variegatum* and *R. microplus* infestations separately, indicating potential species-specific differences in acaricide sensitivity or application efficacy. Another factor could be the widespread use of manual pumps to spread the acaricide preparation on cattle, where ticks can easily be missed. This phenomenon is especially evident in the *Amblyomma variegatum* species, which demonstrates a marked preference for attachment to the lower abdomen (Barré, 1989; Stachurski, 2000), while the *Rhipicephalus microplus* tick is frequently observed on the flanks of animals that are more exposed to acaricides. The alternation of acaricide active substances did not show significant effect on tick infestation at the cattle level. However, its interaction with time since the previous treatment was significant, indicating that the combination of timing and variation in treatment strategy plays a role in controlling tick burden. When excluding the variable “time since the previous acaricide treatment” from the analysis, both the island of origin and cattle breed were significantly associated with tick infestation levels. Cattle from Grande-Terre were less infested than those from Basse-Terre, while no significant difference was observed between cattle from Basse-Terre and Marie-Galante. The more humid climate of Basse-Terre favours tick survival and reproduction (Oshiro et al., 2021; Rahajarison et al., 2014; Solomon and Kaaya, 1998), which likely explains these differences. While Marie-Galante and Grande-Terre share similar climatic conditions, socio-economic factors (Reif et al., 2022) may account for the disparities in infestation levels. Cattle breed also had a significant effect on tick infestation. Crossbred cattle were less infested than Creole cattle, a result that contrasts with previous studies suggesting greater tick resistance in Creole breed (Ben-Jemaa et al., 2023; Camus, 1989; Mollong et al., 2025). This unexpected result may reflect differences in management practices: farmers may provide more attentive care and more frequent acaricide treatments to crossbred animals, which are perceived as more susceptible and represent a greater financial investment due to their genetic value and market potential (Camus, 1989; Shyma et al., 2015). When examining infestations by tick species individually, age was significantly associated with higher infestation levels for both *A. variegatum* and *R. microplus*. Furthermore, male cattle were more heavily infested with the *A. variegatum* tick than females. This sex-based difference in infestation levels aligns with findings by Yessinou et al., 2018, who proposed that male produce more CO2 due to their larger body mass, potentially attracting more ticks over a wider area.

## 6 Conclusion

This study successfully investigated key risk factors associated with tick infestations and *Ehrlichia ruminantium* serology in cattle farms across Guadeloupe. Among the sampled farms, 64% had at least one animal that reacted positively to map1B ELISA. The use of green fodder was significantly associated with increased odds of *E. ruminantium* seropositivity. However, this practice was also linked to reduced tick infestations, indicating the need for further research to clarify the underlying mechanisms. Tick infestation prevalence was high, with 93% of farms and 81% of cattle found to be infested, indicating a substantial increase compared to studies carried in 2003 and 2005. This rise could be attributed to the lack of a coordinated territorial control program for tick management in Guadeloupe’s cattle herds for over a decade. Since 2020, SANIGWA, in collaboration with research institution, has initiated projects aimed at addressing this issue. Tick infestations’ rates were primarily influenced by acaricide use on the territory, and our findings suggest potential emerging resistance to those products, underscoring the urgent need for focussed studies on acaricide resistance in local tick population. Overall, tick and heartwater disease remain major hurdle for breeding activities in Guadeloupe. Reactivating a dedicated surveillance system, such as the previously operational RESPANG plan, would greatly help in gathering essential data to develop effective strategies for managing tick infestations, heartwater, and other tick-borne diseases threatening the livestock sector in the region.

## 7 Acknowledgement

The authors would like to express their gratitude to all the farmers who participated in this study. We would like to acknowledge also the contribution of the colleagues from the TISARU project led by Sylvie Chaumien-Lecollinet (CIRAD) and Jean-Christophe Bambou (INRAE) in the shared survey protocol.

## References

Agreste, 2021. Memento de la statistique agricole - 2021 - Guadeloupe.

Allsopp, B.A., 2015. Heartwater – Ehrlichia ruminantium infection:-EN--FR-La cowdriose – Infection par Ehrlichia ruminantium-ES-Cowdriosis – Infección por Ehrlichia ruminantium. Rev. Sci. Tech. OIE 34, 557–568. 10.20506/rst.34.2.2379

Asamoah, J.K.K., Fatmawati, 2023. A fractional mathematical model of heartwater transmission dynamics considering nymph and adult amblyomma ticks. Chaos, Solitons & Fractals 174, 113905. 10.1016/j.chaos.2023.113905

Baguley, T., 2018. Serious Stat□: A guide to advanced statistics for the behavioral sciences., Bloomsbury publishing. ed.

Barré, N., 1989. Biologie et écologie de la tique Amblyomma Variegatum (Acarina□: Ixodina) en Guadeloupe (Antilles françaises). Université Paris-Sud, France.

Barré, N., Camus, E., Fifi, J., Fourgeaud, P., Numa, G., Rose□Rosette, F., Borel, H., 1996. Tropical Bont Tick Eradication Campaign in the French Antilles Current Status. Annals of the New York Academy of Sciences 791, 64–76. 10.1111/j.1749-6632.1996.tb53512.x

Barré, N., Fargetton, M., Aprelon, R., Coulibandu, L., 1995. Acaricides utilisables dans la lutte contre les tiques aux Antilles□: résultats d’essais en Guadeloupe. Rev. Elev. Med. Vet. Pays Trop. 48, 351–356. 10.19182/remvt.9441

Barre, N., Garris, G.I., Borel, G., Camus, E., 1988. Hosts and Population Dynamics of Amblyomma variegatum (Acari: Ixodidae) on Guadeloupe, French West Indies. Journal of Medical Entomology 25, 111–115. 10.1093/jmedent/25.2.111

Bates, D., Mächler, M., Bolker, B., Walker, S., 2015. Fitting Linear Mixed-Effects Models Using lme4. J. Stat. Soft. 67. 10.18637/jss.v067.i01

Beati, L., Patel, J., Lucas-Williams, H., Adakal, H., Kanduma, E.G., Tembo-Mwase, E., Krecek, R., Mertins, J.W., Alfred, J.T., Kelly, S., Kelly, P., 2012. Phylogeography and Demographic History of Amblyomma variegatum (Fabricius) (Acari: Ixodidae), the Tropical Bont Tick. Vector-Borne and Zoonotic Diseases 12, 514–525. 10.1089/vbz.2011.0859

Ben-Jemaa, S., Adam, G., Boussaha, M., Bardou, P., Klopp, C., Mandonnet, N., Naves, M., 2023. Whole genome sequencing reveals signals of adaptive admixture in Creole cattle. Sci Rep 13, 12155. 10.1038/s41598-023-38774-7

Bernier, R., Yenkamala, L., Philibert, L., Moutoussamy, M., Marie-Magdeleine, C., 2025. Towards plant-based control of Amblyomma variegatum and Rhipicephalus microplus ticks: Knowledge, perceptions, and practices of ruminant livestock farmers in Guadeloupean islands. Veterinary Parasitology: Regional Studies and Reports 60, 101249. 10.1016/j.vprsr.2025.101249

Bianchi, M.W., Barré, N., Messad, S., 2003. Factors related to cattle infestation level and resistance to acaricides in Boophilus microplus tick populations in New Caledonia. Veterinary Parasitology.

Biguezoton, A., Noel, V., Adehan, S., Adakal, H., Dayo, G.-K., Zoungrana, S., Farougou, S., Chevillon, C., 2016. Ehrlichia ruminantium infects Rhipicephalus microplus in West Africa. Parasites & Vectors 9. 10.1186/s13071-016-1651-x

Bolker, B.M., Brooks, M.E., Clark, C.J., Geange, S.W., Poulsen, J.R., Stevens, M.H.H., White, J.-S.S., 2009. Generalized linear mixed models: a practical guide for ecology and evolution. Trends in Ecology & Evolution 24, 127–135. 10.1016/j.tree.2008.10.008

Boucher, F., Moutroifi, Y., Peba, B., Ali, M., Moindjie, Y., Ruget, A.-S., Abdouroihamane, S., Madi Kassim, A., Soulé, M., Charafouddine, O., Cêtre-Sossah, C., Cardinale, E., 2020. Tick-borne diseases in the Union of the Comoros are a hindrance to livestock development: Circulation and associated risk factors. Ticks and Tick-borne Diseases 11, 101283. 10.1016/j.ttbdis.2019.101283

Camus, E., 1989. Etude épidemiologique de la cowdriose a Cowdria ruminantium en Guadeloupe, Etudes et synthèse de l’I.E.M.V.T. IEMVT.

Camus, E., Barre, N., 1995. Vector situation of tick-borne diseases in the Caribbean islands. Veterinary Parasitology 57, 167–176. 10.1016/0304-4017(94)03118-G

Camus, E., Barre, N., 1987. EPIDEMIOLOGY OF HEARTWATER IN GUADELOUPE AND IN THE CARIBBEAN. Onderstepoort Journal of Veterinary Research 54, 419–426.

Core Team, R., 2023. R: A language and environment for statistical computing. R Foundation for Statistical Computing.

Corn, J.L., Barré, N., Thiebot, B., Creekmore, T.E., Garris, G.I., Nettles, V.F., 1993. Potential Role of Cattle Egrets, Bubulcus ibis (Ciconiformes: Ardeidae), in the Dissemination of Amblyomma variegatum (Acari: Ixodidae) in the Eastern Caribbean. Journal of Medical Entomology 30, 1029–1037. 10.1093/jmedent/30.6.1029

Coxe, S., West, S.G., Aiken, L.S., 2009. The Analysis of Count Data: A Gentle Introduction to Poisson Regression and Its Alternatives. Journal of Personality Assessment 91, 121–136. 10.1080/00223890802634175

Dobson, A.J., 2002. An introduction to generalized linear models, 2. ed. ed, Texts in statistical science. Chapman & Hall / CRC, Boca Raton, Fla.

Dufleit V., Guerrini L., Dhune M., Viry L., Belfort A., Jacquet-Crétides L., Deddy J., Cozier J., Naves M., Arthein L., Samut T., Pezeron M., Rodrigues V., Meyer F. D., Etter E., 2025. Prevalence of heartwater in Guadeloupe (2024): stable endemicity and evidence of spread to Les Saintes. bioRxiv 2025.06.11.659135; doi: 10.1101/2025.06.11.659135

Esemu, S.N., Ndip, R.N., Ndip, L.M., 2018. Detection of Ehrlichia ruminantium infection in cattle in Cameroon. BMC Res Notes 11, 388. 10.1186/s13104-018-3479-2

EstradaLPeña, A., Acedo, C.S., Quílez, J., Del Cacho, E., 2005. A retrospective study of climatic suitability for the tick Rhipicephalus (Boophilus) microplus in the Americas. Global Ecology and Biogeography 14, 565–573. 10.1111/j.1466-822X.2005.00185.x

Estrada-Peña, A., Pegram, R.G., Barré, N., Venzal, J.M., 2007. Using invaded range data to model the climate suitability for Amblyomma variegatum (Acari: Ixodidae) in the New World. Exp Appl Acarol 41, 203–214. 10.1007/s10493-007-9050-9

Fabbro, S.D., 2015. Fencing and mowing as effective methods for reducing tick abundance on very small, infested plots. Ticks and Tick-borne Diseases 6, 167–172. 10.1016/j.ttbdis.2014.11.009

Fox, J., Weisberg, S., 2019. An R Companion to Applied Regression, 3rd ed. Thousand Oaks {CA}.

Galan, F., Julien, L., Duflot, B., 2008. Panorama filières animales et typologie systèmes Guadeloupe.

Guo, H., Yin, C., Galon, E.M., Du, J., Gao, Y., Adjou Moumouni, P.F., Liu, M., Efstratiou, A., Lee, S.-H., Li, J., Ringo, A.E., Wang, G., Li, Y., Tumwebaze, M.A., Xuan, X., 2018. Molecular survey and characterization of Theileria annulata and Ehrlichia ruminantium in cattle from Northwest China. Parasitology International 67, 679–683. 10.1016/j.parint.2018.06.011

Hüe, T., 2022. Approche intégrée de la lutte contre la tique du bétail, Rhipicephalus (Boophilus) australis, en Nouvelle-Calédonie (Médecine vétérinaire et santé animale). Université de Nouvelle-Calédonie.

Insee, 2020. ASSOCIATION POUR LA PROTECTION SANITAIRE DES ELEVAGES DE GWADLOUP (SANIGWA) [WWW Document]. https://annuaire-entreprises.data.gouv.fr/. URL https://annuaire-entreprises.data.gouv.fr/entreprise/association-pour-la-protection-sanitaire-des-elevages-de-gwadloup-sanigwa-880672639

Jain, P., Satapathy, T., Pandey, R.K., 2020. Rhipicephalus microplus: A parasite threatening cattle health and consequences of herbal acaricides for upliftment of livelihood of cattle rearing communities in Chhattisgarh. Biocatalysis and Agricultural Biotechnology 26, 101611. 10.1016/j.bcab.2020.101611

Josse, J., Husson, F., 2016. missMDA5□: A Package for Handling Missing Values in Multivariate Data Analysis. J. Stat. Soft. 70. 10.18637/jss.v070.i01

Kerr, S., Greenland, S., Jeffrey, K., Millington, T., Bedston, S., Ritchie, L., Simpson, C.R., Fagbamigbe, A.F., Kurdi, A., Robertson, C., Sheikh, A., Rudan, I., 2023. Understanding and reporting odds ratios as rate-ratio estimates in case-control studies. J Glob Health 13. 10.7189/jogh.13.04101

Laurent, M., Gerbier, G., Faverjon, C., Teissier, R., Redon, J.-M., Vachiery, N., Lefrançois, T., Pradel, J., 2012. Heartwater surveillance in Guadeloupe: a model of partnership between research and surveillance for the Caribbean. Presented at the International Symposium on Veterinary Epidemiology and Economics, Wageningen Academic Publishers, public, pp. 446–446, 1 poster.

Lê, S., Josse, J., Husson, F., 2008. FactoMineR□: An R Package for Multivariate Analysis. J. Stat. Soft. 25. 10.18637/jss.v025.i01

Mapham, D.P.H., Craigie, Vorster, D.J.H., 2017. Heartwater in Cattle and Small Ruminants. On-line Vets (Vets360) 21, 5–11.

Matos, C.A., Gonçalves, L.R., De Souza Ramos, I.A., Mendes, N.S., Zanatto, D.C.S., André, M.R., Machado, R.Z., 2019. Molecular detection and characterization of Ehrlichia ruminantium from cattle in Mozambique. Acta Tropica 191, 198–203. 10.1016/j.actatropica.2019.01.007

Mdladla, K., Dzomba, E.F., Muchadeyi, F.C., 2016. Seroprevalence of Ehrlichia ruminantium antibodies and its associated risk factors in indigenous goats of South Africa. Preventive Veterinary Medicine 125, 99–105. 10.1016/j.prevetmed.2016.01.014

Molia, S., Frebling, M., Vachiéry, N., Pinarello, V., Petitclerc, M., Rousteau, A., Martinez, D., Lefrançois, T., 2008. Amblyomma variegatum in cattle in Marie Galante, French Antilles: Prevalence, control measures, and infection by Ehrlichia ruminantium. Veterinary Parasitology 153, 338–346. 10.1016/j.vetpar.2008.01.046

Mollong, E., Lébri, M., Marie-Magdeleine, C., Lagou, S.M., Naves, M., Bambou, J.-C., 2025. Sustainable management of tick infestations in cattle: a tropical perspective. Parasites Vectors 18. 10.1186/s13071-025-06684-4

Mondry, R., Martinez, D., Camus, E., Liebisch, A., Katz, J.B., Dewald, R., Van Vliet, A.H.M., Jongejan, F., 1998. Validation and Comparison of Three EnzymeLlinked Immunosorbent Assays for the Detection of Antibodies to Cowdria ruminantium Infectiona. Annals of the New York Academy of Sciences 849, 262–272. 10.1111/j.1749-6632.1998.tb11058.x

Mori, Y., Kuroda, M., Makino, N., 2016. Multiple Correspondence Analysis, in: Nonlinear Principal Component Analysis and Its Applications. Springer Singapore, Singapore, pp. 21–28. 10.1007/978-981-10-0159-8_3

Mukhebi, A.W., Chamboko, T., O’Callaghan, C.J., Peter, T.F., Kruska, R.L., Medley, G.F., Mahan, S.M., Perry, B.D., 1999. An assessment of the economic impact of heartwater (Cowdria ruminantium infection) and its control in Zimbabwe. Preventive Veterinary Medicine 39, 173–189. 10.1016/S0167-5877(98)00143-3

Mureithi, D.K., Mukiria, E.W., 2015. An Assessment of Tick Borne Diseases Constraints to Livestock Production in a Smallholder Livestock Production System: a Case of Nijru District, Kenya. International Journal of Research in Agriculture and Forestry 2, 43–49.

Newbold, T., 2023. StatisticalModels [WWW Document]. github.

Oshiro, L.M., Da Silva Rodrigues, V., Garcia, M.V., De Oliveira Souza Higa, L., Suzin, A., Barros, J.C., Andreotti, R., 2021. Effect of low temperature and relative humidity on reproduction and survival of the tick Rhipicephalus microplus. Exp Appl Acarol 83, 95–106. 10.1007/s10493-020-00576-1

Pegram, R., Indar, L., Eddi, C., George, J., 2004. The Caribbean Amblyomma Program: Some Ecologic Factors Affecting Its Success. Annals of the New York Academy of Sciences 1026, 302–311. 10.1196/annals.1307.056

Pegram, R.G., De Castro, J.J., Wilson, D.D., 1998. The CARICOM/FAO/IICA Caribbean Amblyomma Programa. Annals of the New York Academy of Sciences 849, 343–348. 10.1111/j.1749-6632.1998.tb11068.x

Pegram, R.G., Eddy, C., 2003. Progress towards the eradication of Amblyomma variegatum from the Caribbean. Experimental and Applied Acarology 28, 273–281.

Pegram, R.G., Wilsmore, A.J., Lockhart, C., Pacer, R.E., Eddi, C.S., 2007. The Carribean Amblyomma variegatum Eradication Programme: Success or Failure?, in: Vreysen, M.J.B., Robinson, A.S., Hendrichs, J. (Eds.), Area-Wide Control of Insect Pests. Springer Netherlands, Dordrecht, pp. 709–720.

Perreau, P., Morel, P.C., Barre, N., Durand, P., 1980. Existence de la cowdriose (heartwater) à Cowdria ruminantium chez les ruminants des Antilles françaises (La Guadeloupe) et des Mascareignes (La Réunion et Ile Maurice). Revue d’élevage et de médecine vétérinaire des pays tropicaux 33, 21–22.

Peter, S.G., Gakuya, D.W., Maingi, N., Mulei, C.M., 2019. Prevalence and risk factors associated with Ehrlichia infections in smallholder dairy cattle in Nairobi City County, Kenya. Vet World 12, 1599–1607. 10.14202/vetworld.2019.1599-1607

Rahajarison, P., Arimanana, A.H., Raliniaina, M., Stachurski, F., 2014. Survival and moulting of Amblyomma variegatum nymphs under cold conditions of the Malagasy highlands. Infection, Genetics and Evolution 28, 666–675. 10.1016/j.meegid.2014.06.022

Reif, J.-B., Blanc, S., Insee, 2022. Marie-Galante□: une communauté confrontée à un fort repli démographique [WWW Document]. Institut national de statistique et des études économiques.

Rodrigues, V., 2025. WOAH Reference Laboratory Reports activities 2024.

Salas, M., Planchenault, D., Roy, F., 1986. Etude des systèmes d’élevage bovin traditionnel en Guadeloupe. Typologie d’élevage. Revue d’élevage et de médecine vétérinaire des pays tropicaux 39, 59–71.

Semu, S.M., Peter, T.F., Mukwedeya, D., Barbet, A.F., Jongejan, F., Mahan, S.M., 2001. Antibody Responses to MAP 1B and Other Cowdria ruminantium Antigens Are Down Regulated in Cattle Challenged with Tick-Transmitted Heartwater. Clin Diagn Lab Immunol 8, 388–396. 10.1128/CDLI.8.2.388-396.2001

Shyma, K.P., Gupta, J.P., Singh, V., 2015. Breeding strategies for tick resistance in tropical cattle: a sustainable approach for tick control. J Parasit Dis 39, 1–6. 10.1007/s12639-013-0294-5

Solomon, G., Kaaya, G.P., 1998. Development, reproductive capacity and survival of Amblyomma Õariegatum and Boophilus decoloratus in relation to host resistance and climatic factors under field conditions. Veterinary Parasitology 75.

Some, M.V., Biguezoton, A.S., Githaka, N., Adakal, H., Dayo, G.-K., Belem, A., Zoungrana, S., Stachurski, F., Chevillon, C., 2023. The potential of Rhipicephalus microplus as a vector of Ehrlichia ruminantium in West Africa. Ticks and Tick-borne Diseases 14, 102117. 10.1016/j.ttbdis.2022.102117

Spickett, A.M., Heyne, I.H., Williams, R., Spickett, A., 2011. Survey of the livestock ticks of the North West province, South Africa. Onderstepoort Journal of Veterinary Research 78, Art. #305, 12 pages. 10.4102/ojvr.v78i1.305

Stachurski, F., 2000. Invasion of West African cattle by the tick Amblyomma variegatum. Medical Vet Entomology 14, 391–399. 10.1046/j.1365-2915.2000.00246.x

Stachurski, F., Gueye, A., Vachiéry, N., 2019. Cowdriosis/Heartwater, in: Kardjadj, M., Diallo, A., Lancelot, R. (Eds.), Transboundary Animal Diseases in Sahelian Africa and Connected Regions. Springer International Publishing, Cham, pp. 459–484. 10.1007/978-3-030-25385-1_22

Vachiéry, N., Jeffery, H., Pegram, R., Aprelon, R., Pinarello, V., Kandassamy, R.L.Y., Raliniaina, M., Molia, S., Savage, H., Alexander, R., Frebling, M., Martinez, D., Lefrançois, T., 2008. Amblyomma variegatum Ticks and Heartwater on Three Caribbean Islands: Tick Infection and Ehrlichia ruminantium Genetic Diversity in Bovine Herds. Annals of the New York Academy of Sciences 1149, 191–195. 10.1196/annals.1428.081

Vachiéry, N., Meyer, D., Marcelino, I., Alves, P., Raliniaina, M., Stachurski, F., Adakal, H., Sheikboudou, C., Aprelon, R., Pinarello, V., al, et, 2010. Vaccinal approach using inactivated vaccine against heartwater and Ehrlichia ruminantium genetic diversity. Advances in Animal Biosciences 1, 389–390. 10.1017/S2040470010000178

Van Den Heever, M.J.J., Lombard, W.A., Bahta, Y.T., Maré, F.A., 2022. The economic impact of heartwater on the South African livestock industry and the need for a new vaccine. Preventive Veterinary Medicine 203, 105634. 10.1016/j.prevetmed.2022.105634

Weglarczyk, S., 2018. Kernel density estimation and its application. ITM Web Conf. 23, 00037. 10.1051/itmconf/20182300037

WOAH, W.O. for A.H., 2018. Heartwater, in: Terrestrial Manual.

Yessinou, R.E., Adoligbe, C., Akpo, Y., Adinci, J., Youssao Abdou Karim, I., Farougou, S., 2018. Sensitivity of Different Cattle Breeds to the Infestation of Cattle Ticks Amblyomma variegatum, Rhipicephalus microplus, and Hyalomma spp. on the Natural Pastures of Opkara Farm, Benin. Journal of Parasitology Research 2018, 1–9. 10.1155/2018/2570940

Zuur, A.F., Ieno, E.N., Elphick, C.S., 2010. A protocol for data exploration to avoid common statistical problems: Data exploration. Methods in Ecology and Evolution 1, 3–14. 10.1111/j.2041-210X.2009.00001.x

