## Appendix 1 to 5 for "Risk factors associated with tick infestation and *Ehrlichia ruminantium* amongst cattle in Guadeloupe, French West Indies"

Supplementary materials:

Appendix. 1: Results of MCA analysis conducted at farm level for *Amblyomma variegatum*

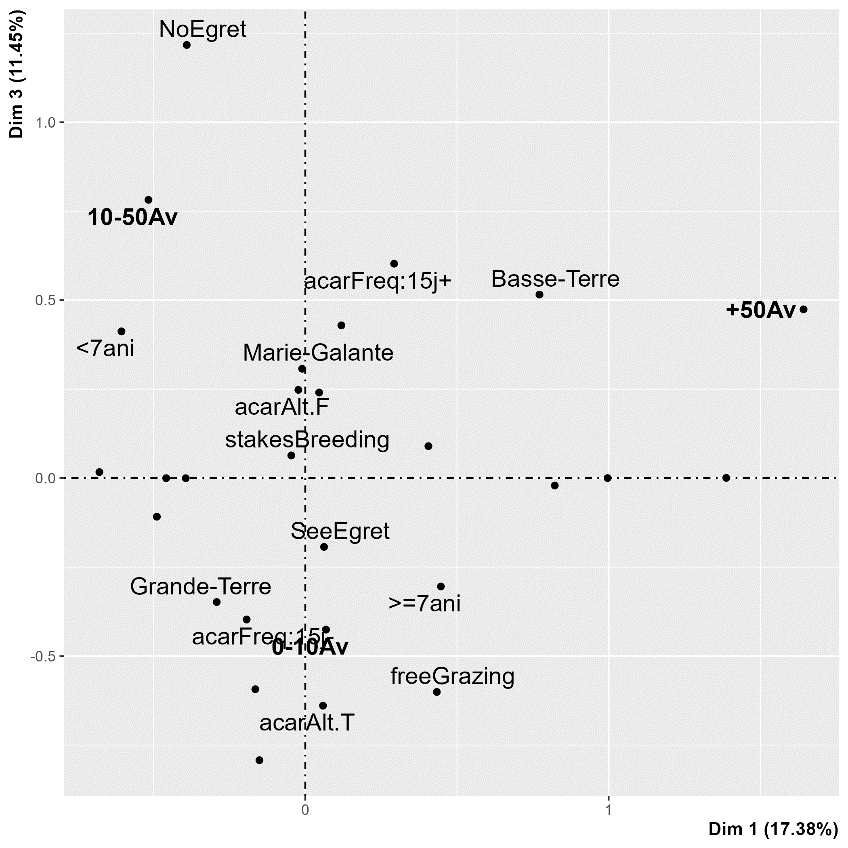

Appendix. 2: Results of MCA analysis conducted at farm level for *Rhipicephalus microplus.*

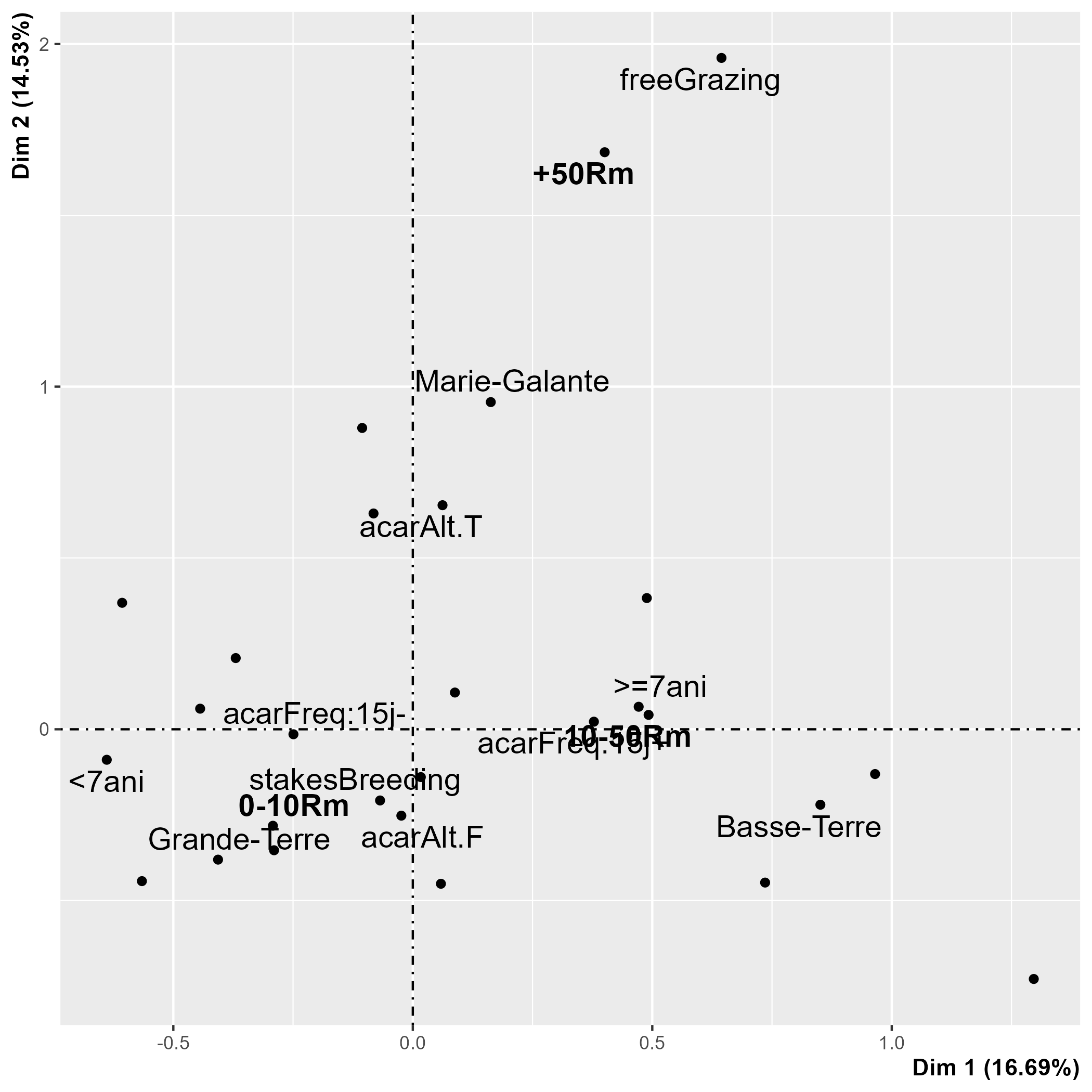

Appendix. 3: Results of MCA analysis conducted at cattle level for *Amblyomma variegatum*.

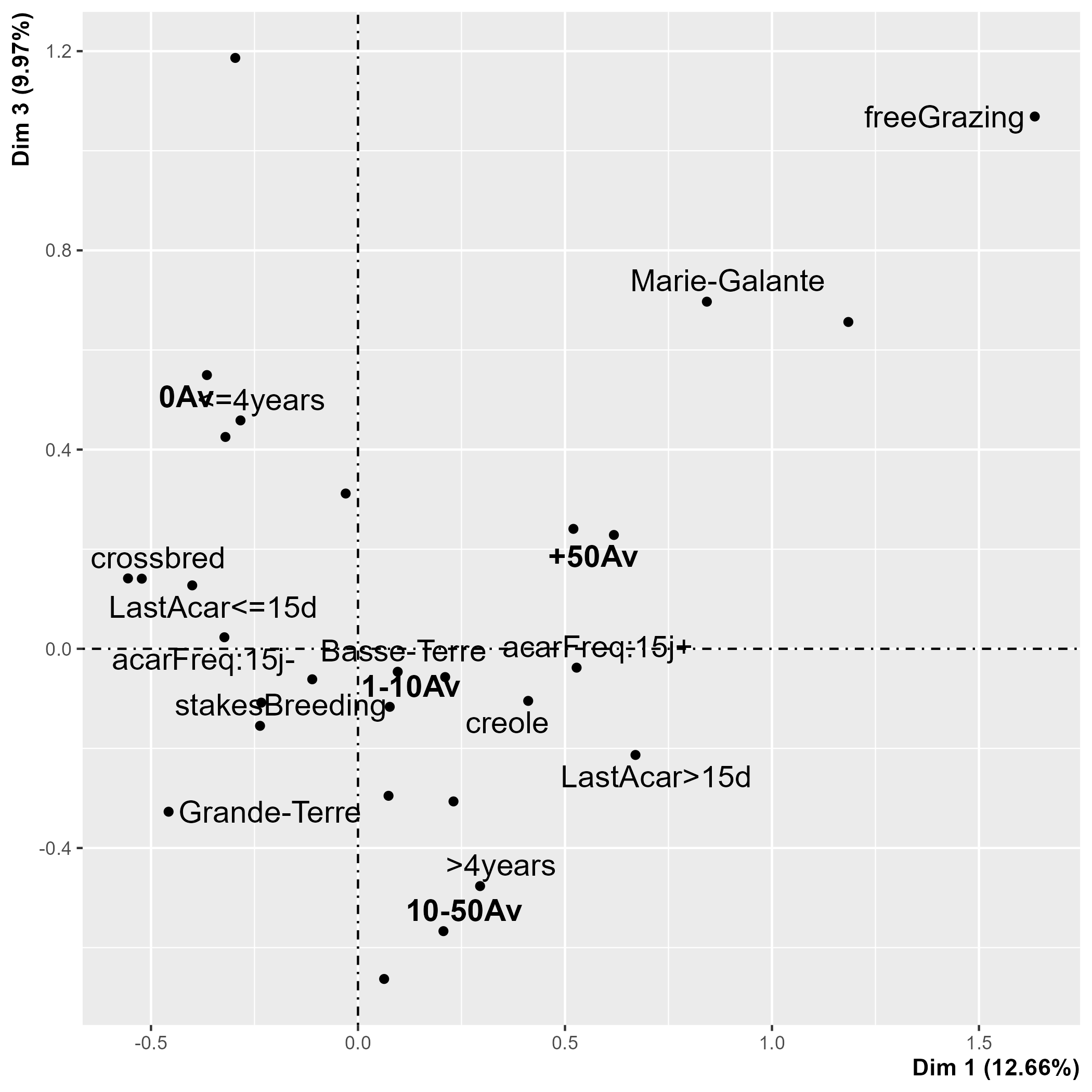

Appendix. 4: Results of MCA analysis conducted at cattle level for *Rhipicephalus microplus*.

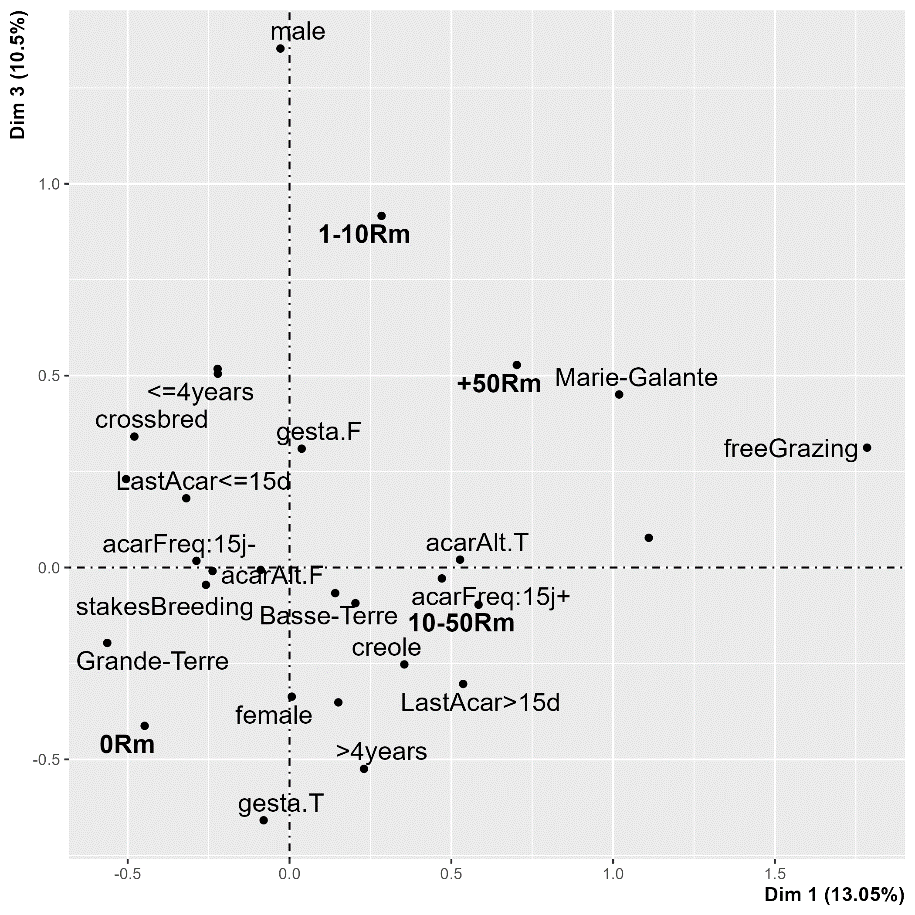

Appendix. 5 : Results of GLM and GLMM analysis conducted at farm and cattle level respectively for *Amblyomma variegatum* and *Rhipicephalus microplus.*

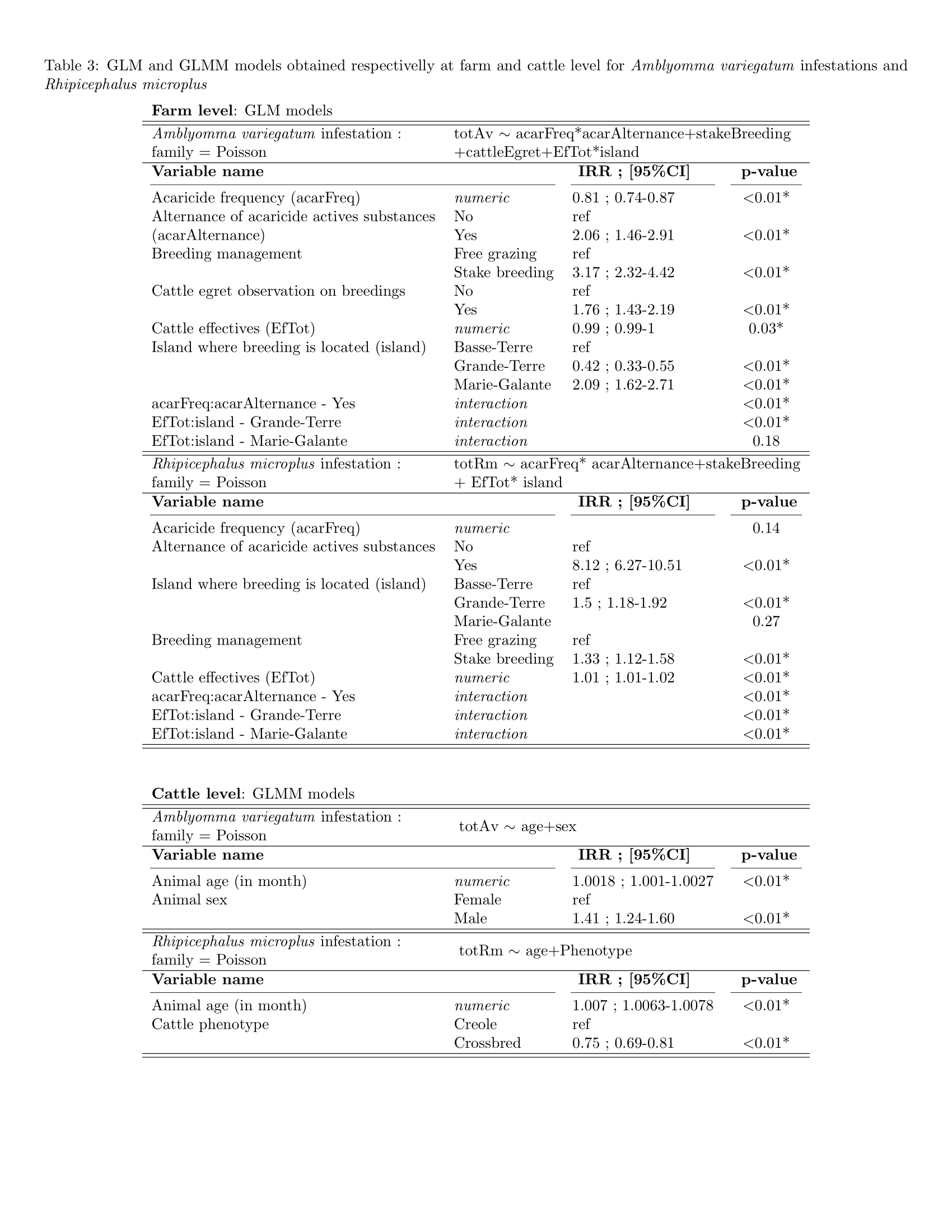
With regard to *Amblyomma variegatum* infestation at farm level, it was found that increasing the frequency of acaricide use significantly reduced *A. variegatum* infestation, with an incidence rate ratio (IRR) of 0.81. There was no effect of acaricide frequency on Rhipicephalus microplus infestations, which may highlight the emergence of acaricide resistance in this tick species. Substance switching significantly increased infestations of *A. variegatum* and *R. microplus*, with IRR of 2.06 and 8.12 respectively. The significant effect of the interaction between acaricide frequency and alternation can be seen in both tick species models. Staking appears to increase tick infestation, with IRRs of 3.17 for *A. variegatum* and 1.33 for *R. microplus*. Island was also found to be a significant factor influencing tick abundance. Marie Galante farms seem to be the most infested with *A. variegatum*, with significant differences from Basse-Terre and an IRR of 2.09. Grande-Terre is significantly less infested than Basse-Terre, with an IRR of 0.42. Interestingly, Grande-Terre is more affected by *R. microplus* than Basse-Terre, with an IRR of 1.5, and no significant difference was found between Marie-Galante and Grande-Terre. The total number of cattle on the farm had an opposite effect on the two species. Increasing the number of cattle reduced the infestation of *A. variegatum*, but seemed to increase the number of *R. microplus*. The interaction between the island and the total number of cattle had a significant effect in the models for both species. Finally, cattle egret observations were found to have a significant positive association with *A. variegatum* tick abundance, potentially highlighting its role as a tick vector in the Caribbean. When considering tick infestation at the individual level, analyses of *A. variegatum* and *R. microplus* revealed a positive effect of age on the abundance of these species, with an IRR slightly greater than one. Male cattle appeared to be significantly more infested with *A. variegatum* than females, and crossbred animals had fewer *Rhipicephalus microplus* than Creole cattle.
